# Serotonergic Modulation of the BNST-CeA Circuit Promotes Sex Differences in Fear Learning

**DOI:** 10.1101/2023.07.03.547577

**Authors:** Rebecca Ravenelle, Carolina Fernandes-Henriques, Jinah Lee, Jia Liu, Ekaterina Likhtik, Nesha S. Burghardt

**Author notes:** Correspondence should be addressed to: Nesha Burghardt Department of Psychology, Hunter College 695 Park Ave, HN 611 New York, NY 10065 Ekaterina Likhtik Department of Biology, Hunter College 695 Park Ave, HN 927 New York, NY 10065.

## Abstract

Post-traumatic stress disorder (PTSD) is characterized by intense fear memory formation and is diagnosed more often in women than men. Here, we show that serotonin differentially affects fear learning and communication in the extended amygdala of male and female mice. Females showed higher sensitivity to the effects of pharmacologically increasing serotonin during auditory fear conditioning, which enhanced fear memory recall in both sexes. Optogenetic stimulation of dorsal raphe terminals in the anterior dorsal bed nucleus of the stria terminalis (adBNST) during fear conditioning increased c-Fos expression in the BNST and central nucleus of the amygdala (CeA), and enhanced fear memory recall via activation of adBNST 5-HT2C receptors in females only. Likewise, in females only, serotonin stimulation during learning enhanced adBNST-CeA high gamma (90-140Hz) synchrony and adBNST-to-CeA communication in high gamma during fear memory recall. We conclude that sex differences in the raphe-BNST-CeA circuit may increase risk of PTSD in women.

## INTRODUCTION

Post-traumatic stress disorder (PTSD) is a stressor and trauma-related disorder that is characterized by intense fearful memory formation. While exposure to severe trauma is a defining feature of PTSD, it is noteworthy that approximately 80% of individuals exposed to a traumatic event do not develop the disorder [1,2]. These findings suggest that in addition to trauma, there are other factors that increase vulnerability to PTSD, one of which is being female. It has been reported that women are at least twice as likely as men to develop PTSD following exposure to trauma [2–4], but the neurobiological factors underlying this sex difference are not known.

Numerous studies implicate serotonin dysfunction in PTSD. For example, PTSD has been associated with polymorphisms in the serotonin-transporter gene [5,6] and a decrease in the density of platelet serotonin-reuptake sites [7–10]. PTSD patients also have an altered response to serotonin agonists [11], including an increase in panic attacks [12]. In rodent studies, enhancing serotonin systemically increases fear conditioning[13–15], a form of associative learning similar to what occurs during the development of PTSD [16]. Although higher levels of serotonin have been reported in female rodents than males [17], the vast majority of serotonin studies with fear conditioning have only been conducted in males and sex differences have not been investigated.

The bed nucleus of the stria terminalis (BNST) is a sexually dimorphic brain region [18–21] that receives dense serotonergic projections from raphe nuclei [22–24] that are activated by foot shocks [25]). While the focus of Pavlovian fear conditioning has historically been on the basolateral amygdala (BLA) [27–29] and the central nucleus of the amygdala (CeA) [30–32], emerging evidence implicates an important role for the BNST in the underlying circuitry. Acquisition of auditory fear conditioning and retrieval of a conditioned fear memory promote activity in the BNST, as measured by upregulation of immediate early gene expression [33,34] and increased firing of BNST neurons [25,35]. Direct stimulation of GABAergic neurons in the BNST during auditory fear conditioning increases tone recall [33], while lesioning the BNST impairs recall of contextual fear [34]. Interestingly, serotonin can modulate fear conditioning through its activity in the BNST [15,25], but again, sex differences have not been explored and downstream areas mediating these effects are not well understood.

The BNST is highly interconnected with the CeA, both of which are components of the extended amygdala. Both regions receive glutamatergic input from the BLA and project to similar brainstem areas that mediate fear responses [36–39]. While the BNST is composed of numerous subdivisions [40–42], the lateral and oval subnuclei, which together comprise the anterolateral BNST, provide the strongest input to the CeA [43]. This input is activated during recall of a contextual fear memory [34], supporting a role for BNST-to-CeA projections in the expression of a conditioned response.

Here, we first tested the effects of acute systemic selective serotonin reuptake inhibitor (SSRI) treatment on auditory fear conditioning in male and female mice. Associated changes in cell activity in the BNST, CeA, and BLA were evaluated with c-Fos immunohistochemistry. We then tested whether there are sex differences in the effects of optogenetically stimulating serotonin in the BNST on fear learning or serotonin receptor expression in this region. Finally, we recorded local field potentials in each sex to investigate how an increase in serotonin in the BNST during learning affects bidirectional communication between the BNST and CeA during retrieval. We found that females were more sensitive than males to SSRI-induced increases in fear learning. Targeted increases of serotonin in the BNST enhanced fear learning in females only, effects attributable to activation of 5-HT_2C_ receptors and the strengthening of input from the adBNST to CeA. These sex specific changes in the raphe-BNST-CeA circuit may underlie sex differences in risk for developing PTSD.

## RESULTS

### Females are more sensitive than males to the effects of acute SSRI treatment on auditory fear conditioning

We first investigated whether stage of estrous cycle influences fear conditioning (Figure S1). Using vaginal cytology, we found that estradiol status on conditioning day did not affect acquisition (group, F_(1,25)_ = 0.14, p = 0.71), tone recall (group, F_(1,25)_ = 0.04, p = 0.84), or recall of contextual fear memory (t_(24)_ = 0.44, p = 0.66) (Figure S1C). Similarly, estradiol status on the day of test did not affect recall of tone (group, F_(1,25)_ = 0.69, p = 0.41) or context (t_(24)_ = 1.56, p = 0.13) (Figure S1D), consistent with rat studies [44,45]. Based on these findings, estrous cycle was not monitored in subsequent experiments.

We then tested the effects of increasing extracellular levels of serotonin throughout the brain on fear conditioning in mice of both sexes. In females, a single injection of the SSRI citalopram (20mg/kg or 10mg/kg, i.p.) before auditory fear conditioning did not affect acquisition (Figure 1B1, group, F_(2,53)_ = 1.24, p = 0.30), but did dose dependently increase tone recall (Figure 1B2, tone x group interaction, F_(8,212)_ = 3.37, p<0.01), with the higher dose increasing freezing during tone-bins 1-3 (20 mg/kg vs. saline, p<0.05) and the lower dose only increasing freezing during tone-bin 1 (10 mg/kg vs. saline, p<0.01). In males, citalopram also increased tone recall (Figure 1C2, tone x group interaction, F_(8,208)_ = 2.63, p<0.01) without affecting acquisition (Figure 1C1, group, F_(2,53)_ = 0.33, p = 0.72), but the effects were not as pronounced as those in females. The lower dose did not affect freezing to any of the tones (10 mg/kg vs. saline, p>0.05) and the higher dose only increased freezing during the first tone-bin (20 mg/kg vs. saline, p<0.05). When recall of contextual fear memory was tested, we found no effect of citalopram in females (Figure 1B3, F_(2,53)_ = 1.62, p = 0.21) or males (Figure 1C3, F_(2,53)_ = 0.86, p = 0.43). Together, our results are consistent with previous work in rats showing that SSRIs enhance tone recall[13–15], but indicate that females are more sensitive than males to these effects.

**Figure 1.**
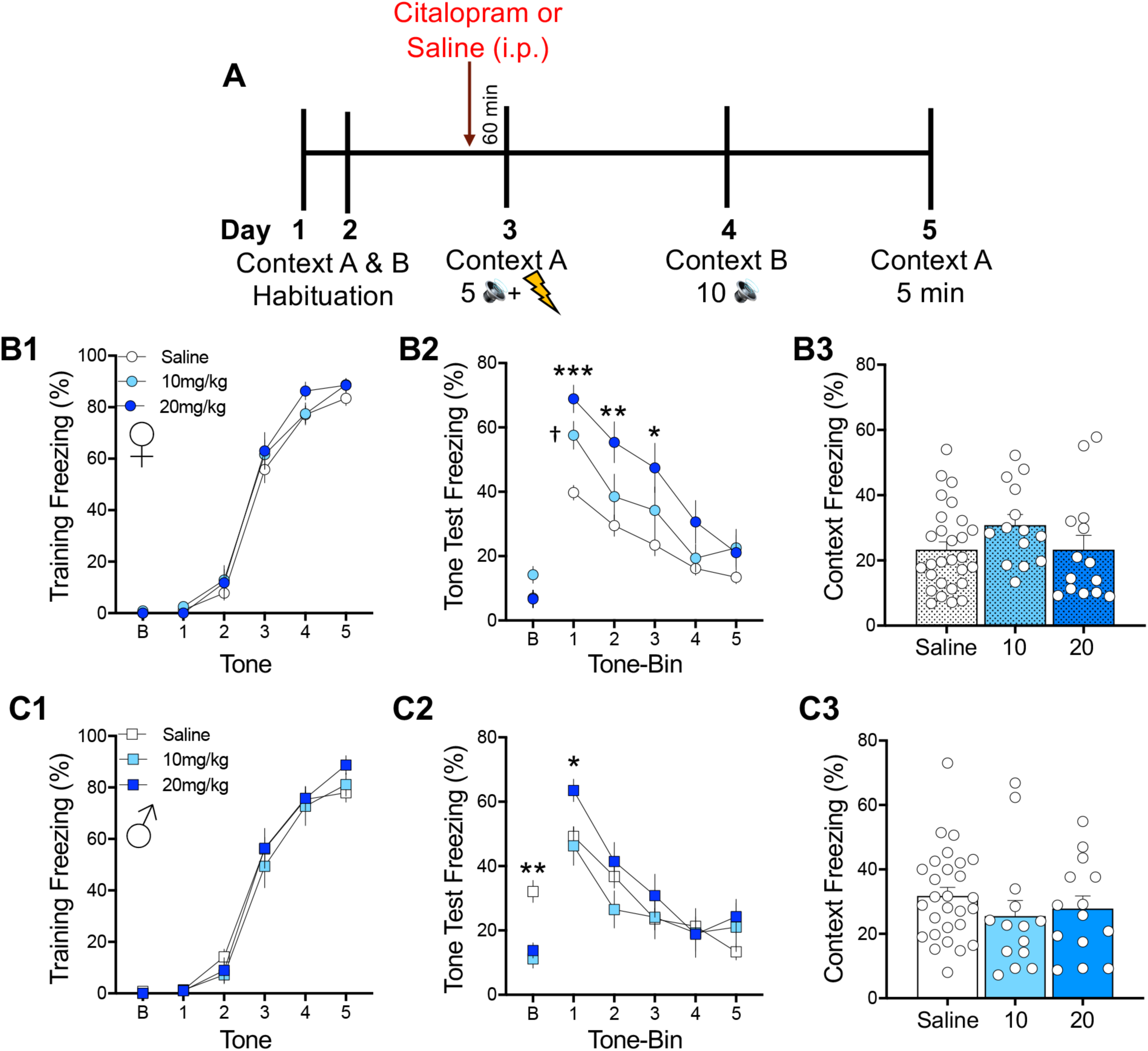
Acute systemic administration of the SSRI citalopram prior to auditory fear conditioning has a stronger effect on tone recall in females than males. **(A)** Schematic of procedures. Mice were habituated to the training (Context A) and testing (Context B) contexts before receiving an i.p. injection of citalopram (10mg/kg or 20mg/kg) or saline 60 minutes before fear conditioning. On the following two days, mice were tested drug-free to 10 presentations of the tone alone (Context B) and to the training context (Context A). **(B-C)** Acute citalopram treatment did not affect percentage of time spent freezing to tones during conditioning in **(B1)** females or **(C1)** males. When mice were tested drug-free to presentations of the tone alone, both citalopram doses increased tone-evoked freezing in **(B2)** females, while **(C2)** males were only affected by the higher dose. **(C2)** In males, citalopram decreased baseline (B) freezing prior to presentation of the first tone on the testing day. Citalopram did not affect recall of contextual fear memory in **(B3)** females or **(C3)** males. n = 14 females per citalopram dose; n = 28 combined saline females; n = 14 males per citalopram dose; n = 28 combined saline males. Data are represented as mean ± SEM. ***p<0.0001 vs. saline, ** p<0.01 vs. saline, * p<0.05 vs. saline, ^†^p<0.01 vs. saline.

Interestingly, in males both doses of citalopram reduced baseline freezing measured 30 seconds before the first tone test (Figure 1C2, Figure S2A, F_(2,52)_ = 12.72, p<0.0001; 10 mg/kg p < 0.001; 20 mg/kg p < 0.01). This effect was investigated further in a separate cohort of males injected with the same two doses of citalopram before auditory fear conditioning and tested in Context B in the absence of tones (Figure S2B). We again found that citalopram did not affect acquisition (Figure S2C1, group, F_(2,22)_ = 0.90, p = 0.42) but did decrease freezing to a different context the next day (Figure S2C2, F_(2,22)_ = 14.14, p <0.001; 10 mg/kg p<0.01; 20 mg/kg p <0.001). However, freezing exhibited by the saline group rapidly declined and reached levels comparable to the citalopram groups after the first two tones are usually presented (time-bin 1) (time-bin x group interaction, F_(8,88)_ = 3.55, p<0.01, Tukey’s HSD, time-bin 2-5, n.s.), indicating that group differences are short lasting. In the absence of tone testing, we again found no effect of citalopram on freezing to the training context (Context A) (Figure S2C3, F_(2,22)_ = 0.39, p = 0.68).

### Acute systemic administration of the lower dose of citalopram prior to auditory fear conditioning increases c-Fos expression in the adBNST and CeA of females only

Our finding that the lower dose of citalopram (10 mg/kg) did not affect acquisition but did promote tone recall when females (but not males) were tested drug-free, indicates that this dose affects memory consolidation in a sex-specific manner. We investigated the neural correlates of this effect by administering citalopram (10mg/kg) before auditory fear conditioning and quantifying training-induced upregulation of c-Fos in the BNST, CeA and BLA of males and females during early consolidation. Within the BNST, we focused on the oval, lateral, and medial subregions, which together comprise the anterior dorsal BNST (adBNST). Citalopram- and saline-treated mice were perfused 90 minutes after fear conditioning (trained) and the number of c-Fos-positive cells was compared to non-trained mice perfused from the home cage (naïve). The time between the injection (citalopram or saline) and perfusion was the same for all groups (160 minutes) (Figure 2A). In the adBNST of females, we found that citalopram increased the number of c-Fos+ cells in the oval (drug x training interaction, F_(1,12)_ = 4.80, p < 0.05; trained drug vs. trained saline, p<0.01) and lateral (drug x training interaction, F_(1,12)_ = 6.53, p < 0.05; trained drug vs. trained saline, p<0.05) subregions of trained mice, with effects that approached significance in the medial subregion (drug x training interaction, F_(1,12)_ = 2.11, p = 0.17, trained drug vs. trained saline, p = 0.07) (Figure 2B, 2C). Citalopram did not affect c-Fos in naive mice (p >0.05). In the absence of drug, conditioning only upregulated c-Fos in the medial subregion (training, F_(1,12)_ = 50.82, p<0.0001; naïve saline vs. trained saline, p <0.01) (Figure 2C). However, in males citalopram had no effect in the adBNST (drug x training interaction, ov: F_(1,12)_ = 0.06; al: F_(1,12)_ = 0.14; am: F_(1,12)_ = 1.24, p>0.05). Similar to females, there was a main effect of training in the medial subregion (F_(1,12)_ = 4.77, p < 0.05) that appeared to be driven by the saline group, but none of the post-hoc comparisons reached significance (Figure 2D).

**Figure 2.**
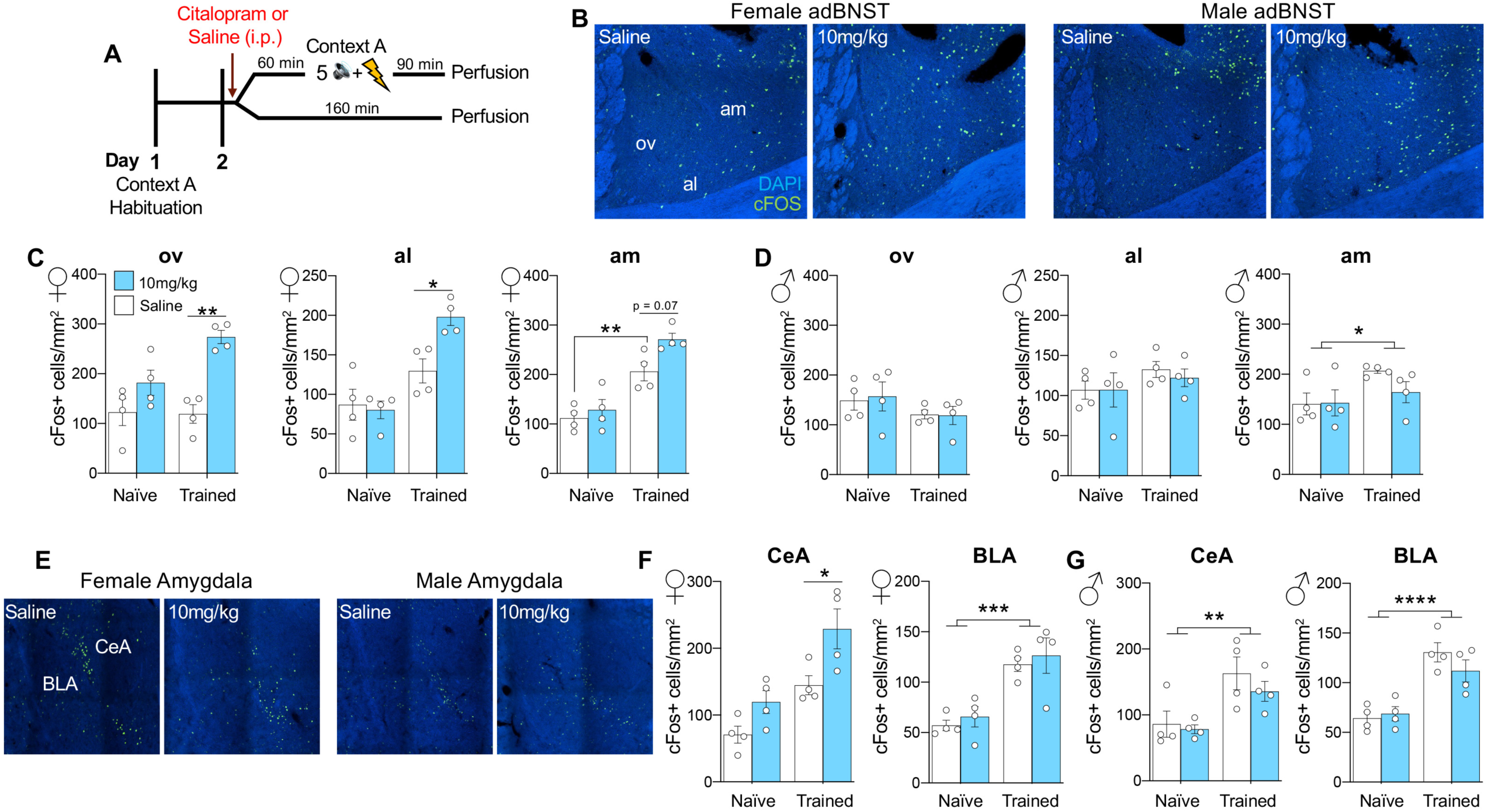
Acute systemic administration of citalopram prior to auditory fear conditioning increases c-Fos expression in the adBNST and CeA of females only. **(A)** Schematic of procedures. Mice were habituated to the training context (Context A) one day before receiving an i.p. injection of citalopram (10mg/kg) or saline. The trained group was fear conditioned 60 minutes after the injection and perfused 90 minutes post-training. The naïve group was perfused 160 minutes after the injection. **(B)** Images of cells expressing c-Fos (green) in the adBNST of trained females (left) and trained males (right). **(C-D)** Quantification of cells expressing c-Fos in the adBNST of **(C)** females and **(D)** males. Citalopram enhanced c-Fos expression in all 3 subregions of the adBNST in trained females only. **(E)** Images of cells expressing c-Fos (green) in the amygdala of trained females (left) and males (right). **(F-G)** Quantification of c-Fos+ cells in the CeA and BLA of **(F)** females and **(G)** males. Citalopram enhanced c-Fos expression in the CeA of trained females only. n = 4 females per group; n = 4 males per group. Data are represented as mean ± SEM. ov = oval BNST, al = lateral BNST, am = medial BNST, CeA = central nucleus of the amygdala, BLA = basolateral amygdala. *p < 0.05, **p < 0.01, ***p<0.001, ****p<0.0001.

In the amygdala of both sexes (Figure 2E-2G), fear conditioning upregulated c-Fos in the BLA (female training, F_(1,12)_ = 30.18, p < 0.001; male training, F_(1,12)_ = 39.27, p < 0.0001), as expected. This was not further augmented by citalopram (effects of drug and interactions, n.s.). In the CeA, fear conditioning also upregulated c-Fos in both sexes (female training, F(_1,12_) = 21.64, p<0.001; male training, F(_1,12_) = 13.98, p<0.01), an effect that was enhanced by citalopram in females but not males (female drug, F_(1,12)_ = 11.43, p<0.01, trained drug vs. trained saline, p<0.05). Together, we found that acute SSRI treatment promotes training-induced upregulation of c-Fos expression in the adBNST and CeA of females, but not males, implicating a role for these regions in mediating sex differences in fear learning when serotonin levels are increased.

### Optogenetic stimulation of 5-HT terminals in the adBNST during auditory fear conditioning enhances tone recall and upregulates c-Fos expression in the oval BNST and CeA of females only

Using the TpH2-ChR2-YFP-Bac mouse line [46], we next tested whether selectively increasing serotonin in the adBNST is sufficient to produce sex differences in fear learning (Figure 3A-3C). We first determined that the number of TpH2-ChR2+ neurons projecting from the dorsal and medial raphe nuclei (DRN and MRN) to the adBNST is similar in males and females in this mouse line (Figure S3, p>0.05). Then we optogenetically stimulated serotonin in the adBNST during auditory fear conditioning by administering blue light during each tone presentation and found no effect of stimulation on acquisition in either sex (Figure 3D1, females, genotype, F_(1,42)_ = 0.03, p = 0.86; Figure 3E1, males, genotype, F_(1,41)_ = 0.63, p = 0.43). The next day, females that had been stimulated exhibited increased tone evoked freezing (Figure 3D2, genotype, F_(1,42)_ = 12.94, p<0.001), but stimulated males did not (Figure 3E2, genotype, F_(1,41)_ = 0.002, p = 0.96). Similar to our findings with citalopram, non-stimulated males froze more than stimulated males at baseline (Figure 3E2, S4A, t_(41)_ = 3.63, p < 0.001), but rapidly decreased freezing to the testing context by the first tone test (Figure S4C). When mice were returned to the training context, there was no effect of previous stimulation on contextual freezing in females (Figure 3D3, t_(42)_ = 0.72, p = 0.48) or males (Figure 3E3, t_(41)_ = 1.51, p = 0.14). Together, these results demonstrate that increasing serotonin in the adBNST enhances tone recall in females only, with no effect on contextual fear memory in either sex.

**Figure 3.**
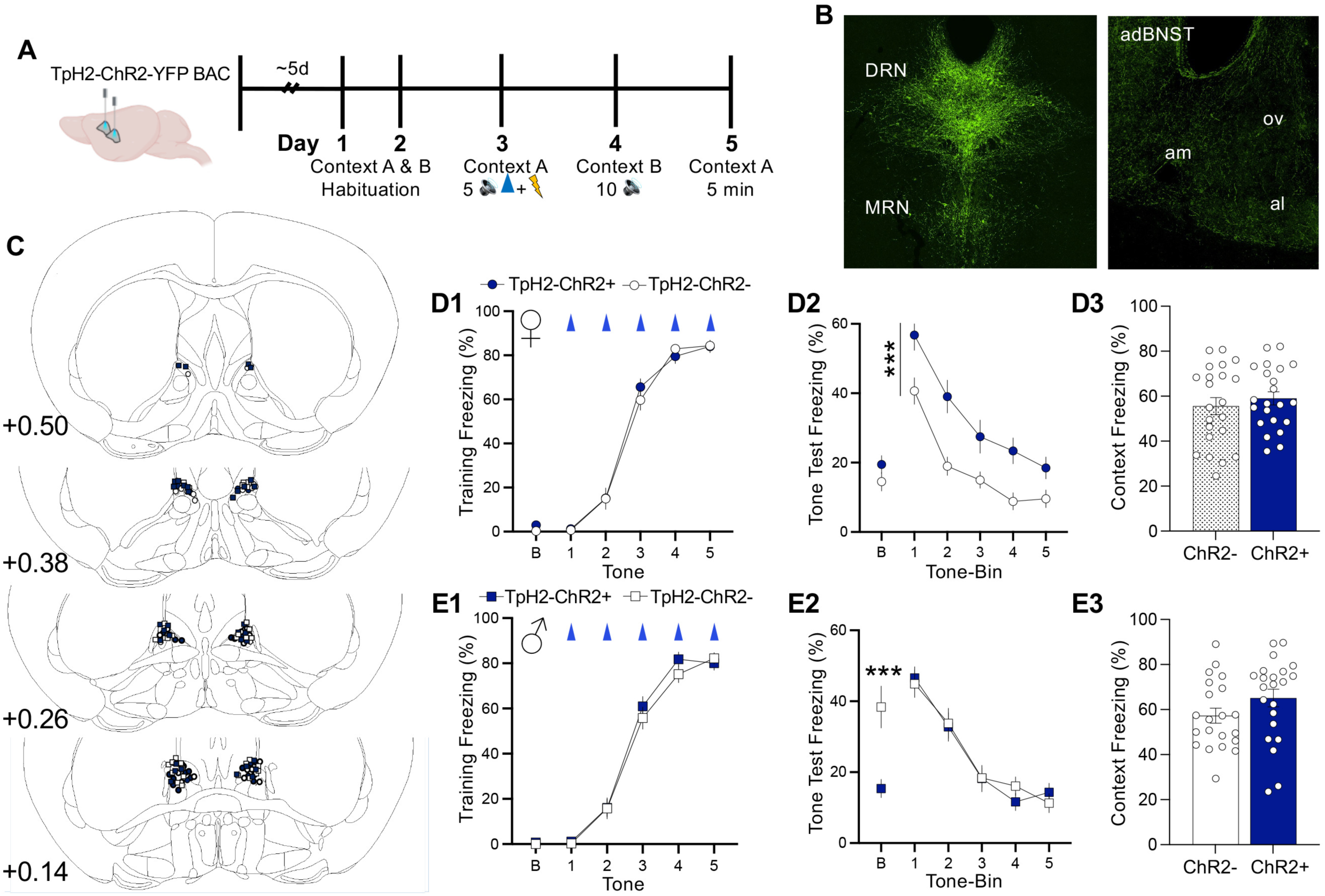
Optogenetic stimulation of 5-HT terminals in the adBNST during auditory fear conditioning enhances tone recall in females only. **(A)** Schematic of procedures. Mice were implanted bilaterally with fiber optics in the adBNST before they were habituated to the training (Context A) and testing (Context B) contexts. During auditory fear conditioning (5 CS-US pairings), a laser delivered blue light to the adBNST during each of the 5 tones. On the following two days, mice were tested to 10 tones (Context B) and the training context (Context A) in the absence of shock or laser stimulation. **(B)** Expression of ChR2-EYFP in the raphe nuclei (left) and adBNST (right). **(C)** Histological verification of optic fiber placements. **(D-E)** Optogenetic stimulation of 5-HT in the adBNST did not affect freezing to the tone during conditioning in **(D1)** females or **(E1)** males. During the tone test, **(D2)** stimulated females but not **(E2)** males exhibited increased tone-evoked freezing. Stimulated males exhibited decreased baseline (B) freezing prior to presentation of the first tone. Optogenetic stimulation did not affect recall of contextual fear memory in **(D3)** females or **(E3)** males. n = 22 females per group; n = 22 ChR2+ males; n = 21 ChR2-males. Data are represented as mean ± SEM. *** p<0.001.

To test whether this sex difference is associated with differential activity in the extended amygdala during learning, we optogenetically stimulated serotonin in the adBNST during fear conditioning and perfused mice 90 minutes later (trained) (Figure 4A-4B). The number of c-Fos+ cells was compared to non-trained mice that were perfused 90 minutes after receiving light stimulation in a familiar cage (naïve). In the adBNST, we found that serotonin stimulation increased the number of c-Fos+ cells in the oval subregion of trained females (Figure 4C, genotype x training interaction, F_(1,12)_ = 6.45, p <0.05); trained ChR2-vs. trained ChR2+, p<0.05), but not trained males (Figure 4D, genotype x training interaction, F_(1,11)_ = 0.66, p = 0.43). While fear conditioning upregulated c-Fos in the lateral (female, training, F_(1,12)_ = 18.43, p < 0.01; male, training, F_(1,11)_ = 9.60, p < 0.05) and medial subregions of both sexes (female, training F_(1,12)_ = 44.73, p < 0.0001; male, training, F_(1,11)_ = 30.81, p < 0.001), this was not further enhanced by stimulation (Figure 4C-4D, interactions n.s.). As with systemic citalopram treatment, selectively increasing serotonin in the adBNST promoted learning-induced upregulation of c-Fos in the CeA of females only (Figure 4E-4G, female genotype x training interaction, F_(1,12)_ = 19.52, p < 0.001, female training, F_(1,12)_ = 100.2; p < 0.0001, naïve ChR2-vs. trained ChR2-, p<0.01, trained ChR2-vs. trained ChR2+, p<0.001; male training, F_(1,11)_ = 45.88, p < 0.0001). Although fear conditioning upregulated c-Fos in the BLA of both sexes (female, training, F_(1,12)_ = 21.58; p < 0.01; male, training, F_(1,11)_ = 32.05; p < 0.001), this was not further increased with optogenetic stimulation (effects of genotype and interactions, n.s.). Together these findings indicate that selectively increasing serotonin in the adBNST during learning increases cell activity in the oval nucleus and promotes activity downstream in the CeA in a sexually dimorphic manner.

**Figure 4.**
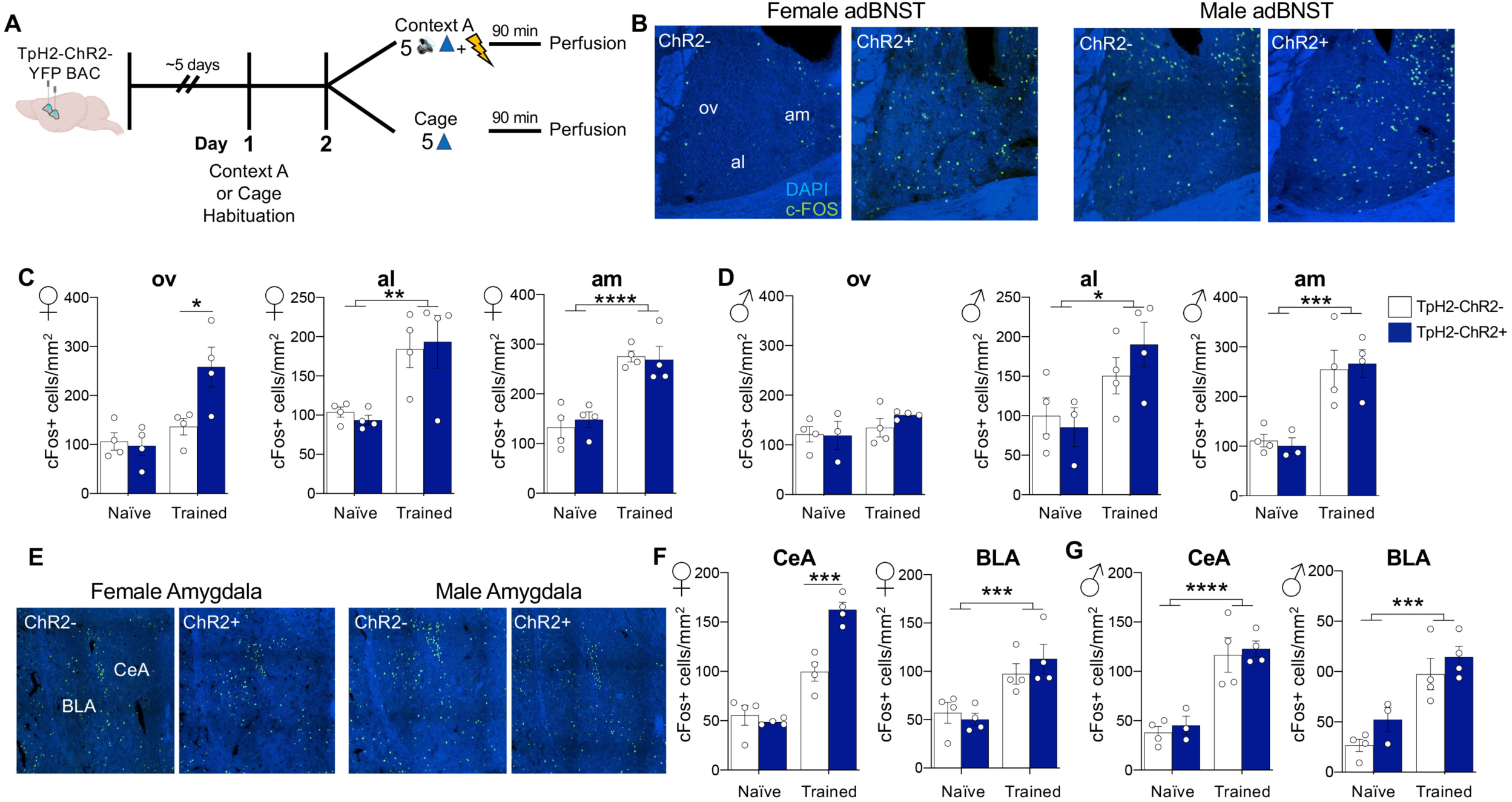
Optogenetic stimulation of 5-HT terminals in the adBNST during auditory fear conditioning increases c-Fos expression in the adBNST and CeA of females only. **(A)** Schematic of procedures. Mice were bilaterally implanted with fiber optics in the adBNST. After recovery, the trained group was habituated to the training context (Context A) the day before being fear conditioned with 5 tone-shock pairings. Blue light was delivered to the adBNST during each tone presentation. The naïve group was habituated to a novel cage the day before they received 5 bouts of blue light in the adBNST in that cage. Both groups were perfused 90 minutes post-stimulation. **(B)** Images of cells expressing c-Fos (green) in the adBNST of trained females (left) and trained males (right). **(C-D)** Quantification of cells expressing c-Fos in the adBNST of **(C)** females and **(D)** males. Optogenetic stimulation only increased c-Fos expression in the oval subregion of trained females. **(E)** Images of c-Fos+ cells in the amygdala of trained females (left) and trained males (right). **(F-G)** Quantification of c-Fos+ cells in the CeA and BLA of **(F)** females and **(G)** males. Optogenetic stimulation of 5-HT terminals in the adBNST enhanced c-Fos expression in the CeA of trained females only. n = 4 females per group; n = 3-4 males per group. Data are represented as mean ± SEM. ov = oval BNST, al = lateral BNST, am = medial BNST, CeA = central nucleus of the amygdala, BLA = basolateral amygdala. *p < 0.05, **p < 0.01, ***p < 0.001, ****p < 0.0001.

### Blocking 5-HT_2C_ receptors in the adBNST prevents 5-HT-induced enhancement of tone recall in females

The BNST is a sexually dimorphic brain region, but it is not known if there are sex differences in serotonin signaling in this area. We used qRT-PCR to investigate whether transcript levels of the 5-HT_1A_ (*Htr1a*), 5-HT_2A_ (*Htr2a*), 5-HT_2C_ (*Htr2c*), 5-HT_3_ (*Htr3a*), and 5-HT_7_ (*Htr7*) receptors in the adBNST are different in males and females. Transcript levels of somatostatin (*Sst*) were also quantified, as this type of BNST neuron is implicated in the regulation of fear [33]. Transcript levels were the highest for *Htr1a* and *Htr2c* in both sexes (Figure 5A-B, females, F_(4,35)_ = 35.25, p < 0.0001; males, F_(4,35)_ = 20.98, p < 0.0001), but the relative expression of *Htr2c* to *Htr1a* was higher in females than males (Figure 5C, t_(14)_ = 2.27, p < 0.05). We also identified a sex difference in 5-HT_2C_ receptors, but not the other serotonin receptors or somatostatin, with females expressing higher *Htr2c* transcript levels than males (Figure 5D-E, *Htr2c*, t_(14)_ = 2.51, p < 0.05; *Htr1a*, t_(14)_ = 1.42, p = 0.18; *Htr2a*, t_(14)_ = 1.10, p = 0.29; *Htr7*, t_(14)_ = 0.29, p = 0.77; *Htr3a*, t_(14)_ = 1.39, p = 0.19; *Sst*, t_(14)_ = 1.04, p = 0.32). To determine whether activation of 5-HT_2C_ receptors in the adBNST contributes to serotonin-induced increases in fear learning in females, we pharmacologically blocked them with a local infusion of a 5-HT_2C_ antagonist (RS102221) 10-15 minutes before females were fear conditioned with optogenetic stimulation of serotonin (Figure 6A). A mixed effects model for repeated measures revealed no effect of the antagonist on acquisition (Figure 6B, tone x group, F_(6,48)_ = 2.26, p = 0.053; group, F_(2,16)_ = 1.25, p = 0.31). However, the antagonist did block serotonin-induced increases in tone recall the next day when mice were tested drug-free without light stimulation (Figure 6C1, tone x group interaction, F_(8,64)_ = 2.09, p<0.05; ChR2+/aCSF vs. ChR2+/antagonist, p<0.05; ChR2-/aCSF vs. ChR2+/antagonist, p>0.05). This was confirmed with a one-way ANOVA on freezing during the first 2 tone bins (tones 1-4) (Figure 6C2, F_(2,16)_ = 4.23, p<0.05; Holm-Sidak’s, p<0.05). When mice were returned to the training context, there were no group differences in contextual freezing (Figure 6D, F_(2,16)_ = 0.96, p = 0.41). To verify that the effect of the 5-HT_2C_ antagonist on tone recall was temporary, a subset of antagonist-treated mice was reconditioned with blue light following a local infusion of aCSF (Figure S5A). Mice froze significantly more during the second conditioning session than the first (Figure S5B, training session x tone interaction, F_(4,36)_ = 7.61, p<0.001). This increase was already detectable at baseline (t_(9)_ = 2.67, p<0.05) and during the first tone presentation (p<0.01), both of which were before administration of the first shock of the reconditioning session, indicating memory of the first training session. Importantly, reconditioning in the absence of antagonist significantly increased tone recall (Figure S5C, test session, F_(1,9)_ = 41.85, p<0.001), demonstrating that the circuits underlying memory retrieval remained intact.

**Figure 5.**
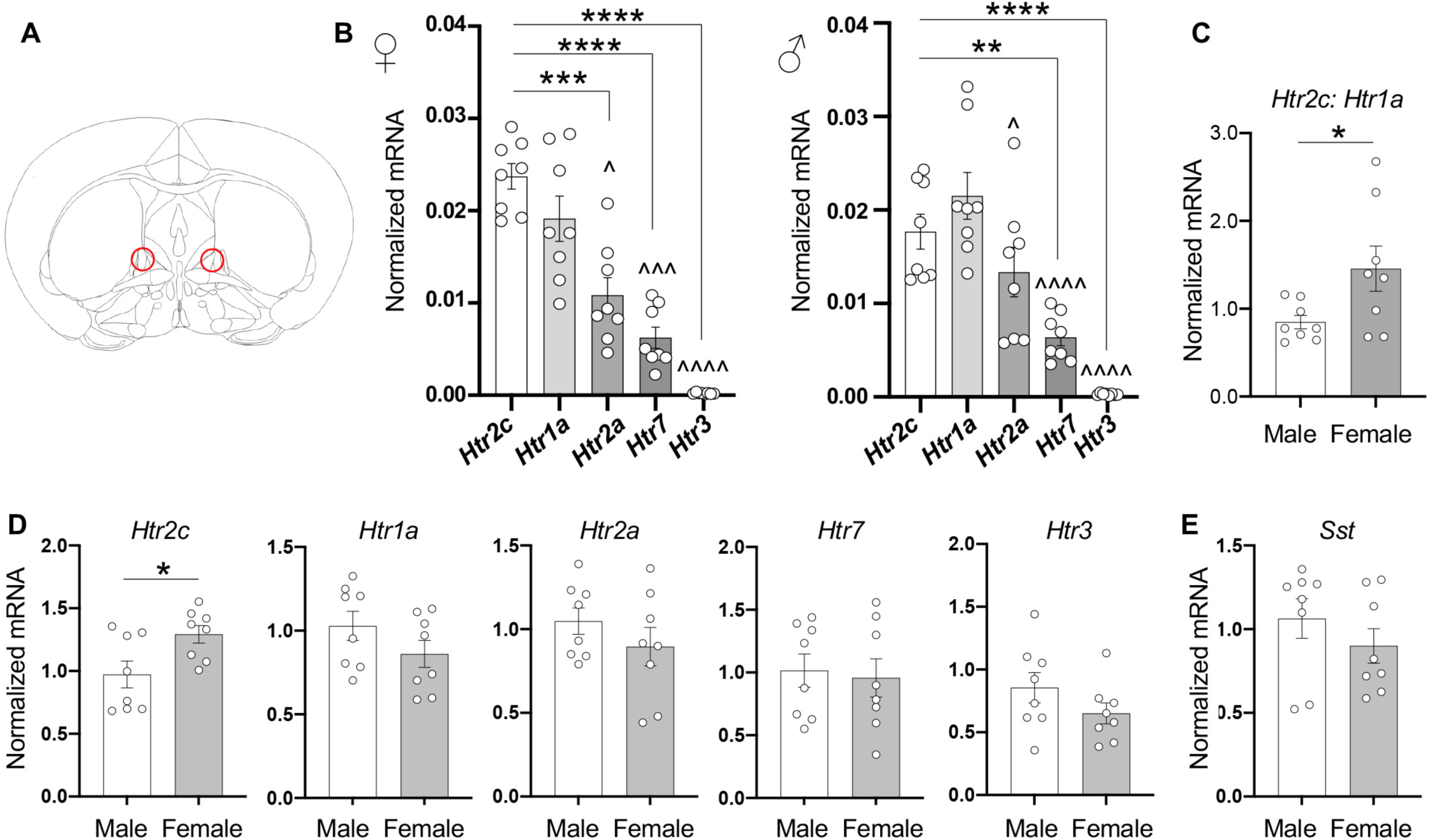
Females express more *Htr2c* transcript than males in the adBNST. **(A)** Location of bilateral adBNST tissue punches (1mm) from TpH2-ChR2-mice for qRT-PCR analysis. **(B)** Transcript levels of the 5-HT_1A_ (*Htr1a*), 5-HT_2A_ (*Htr2a*), 5-HT_2C_ (*Htr2c*), 5-HT_7_ (*Htr7*), and 5-HT_3_ (*Htr3a*) receptors in the adBNST of females (left) and males (right). Data are expressed as transcripts normalized to housekeeping references, which are *Gapdh* and *18s*. In both sexes, expression was the highest for *Htr1a* and *Htr2c*. ^p<0.05 vs. *Htr1a*, ^^^p<0.001 vs. *Htr1a*, ^^^^p<0.0001 vs. *Htr1a*; **p<0.01, ***p<0.001, ****p<0.0001. **(C)** Relative expression of *Htr2C* to *Htr1a* was higher in females than males. **(D-E)** There is a sex difference in *Htr2c* only, with females expressing more transcript than males. Data are expressed as transcripts normalized to male levels. Each data point was acquired from bilateral tissue punches combined from 2 mice (n = 8 combined samples per sex; n = 16 mice per sex). Data are represented as mean sample ± SEM. *p<0.05.

**Figure 6.**
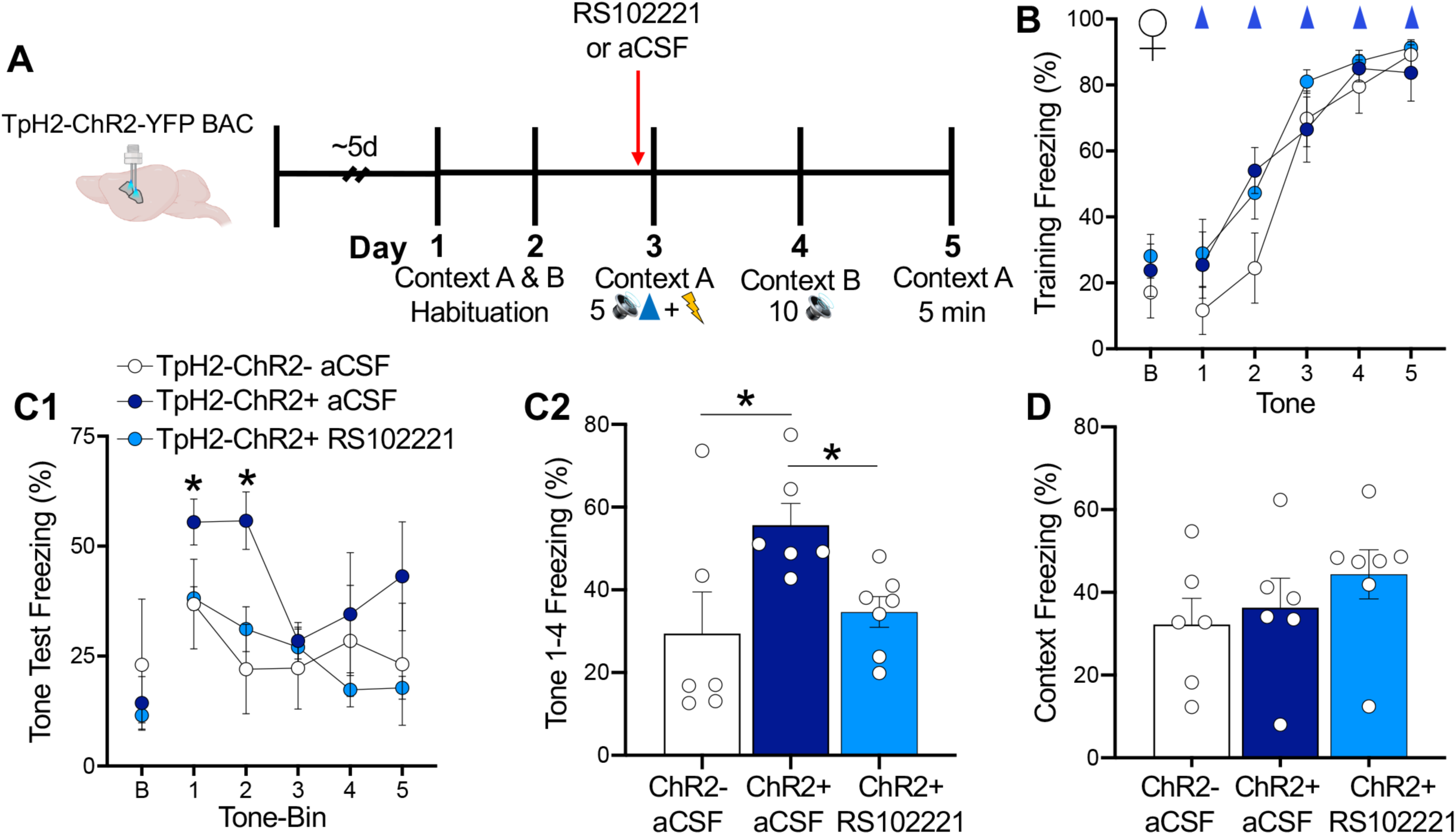
Blocking 5-HT_2C_ receptors in the adBNST prevents 5-HT-induced enhancement of tone recall in females. **(A)** Schematic of procedures. Female mice received a direct bilateral infusion of the 5-HT_2C_ receptor antagonist RS102221 or aCSF into the adBNST 10-15 minutes before fear conditioning. During conditioning, a laser delivered blue light to the adBNST during each of the 5 tones. Mice were tested to the tone and training context on the following two days without laser stimulation, antagonist, or shock. **(B)** Pre-treatment with RS102221 did not affect percentage of time spent freezing to the tone during conditioning. **(C1-C2)** During the tone test, infusion of RS102221 blocked the 5-HT-induced enhancement in freezing to the first 4 tones (tone-bin 1-2; *p < 0.05 ChR2+/aCSF vs. ChR2+/RS102221 and ChR2-/aCSF) and did not affect (**D**) contextual fear memory. n = 6-7 per group. Data are represented as mean ± SEM.

### Inhibition of projections from the CeA to the adBNST impairs recall of tone and context in both sexes

Our c-Fos data demonstrating that increasing serotonin during learning increases activity in the adBNST and CeA in a sex-specific manner prompted us to ask whether there is also a sex difference in communication between these regions. We began investigating this possibility by focusing on output projections from the CeA, a region long known to be required for the acquisition and expression of conditioned fear [30,32,47,48]. Using DREADDs, we tested the effects of inhibiting the CeA-to-adBNST pathway on auditory and contextual fear conditioning in males and females (Figure 7A, 7B, S6B). To account for the possibility that this pathway is modulated by serotonin in females, we optogenetically stimulated serotonin in the adBNST of females during auditory fear conditioning, as above. Based on our behavioral findings indicating no effect of serotonin on this circuit in males, only ChR2-males were tested using the same protocol. We found that inhibiting projections from the CeA to the adBNST with a local infusion of CNO immediately before training impaired acquisition in both sexes (Figure 7C1, mixed effects model, females, virus, F_(1,27)_ = 16.00, p<0.001, interactions, n.s.; Figure 7D1, males, virus, F_(1,14)_ = 17.85, p<0.001). In females, this effect was driven by ChR2+ mice with the DREADD virus (Tone-bin 5: Fisher’s LSD, ChR2+hM4Di vs. all other groups, p<0.01). Interestingly, inhibition during training led to impaired retrieval of the tone and context in females (Figure 7C2, tone recall, tone x genotype x virus, F_(4,112)_ = 2.87, p<0.05; virus, F_(1,28)_ = 13.24, p<0.01; Figure 7C3, context recall, virus, F_(1,28)_ = 17.25, p<0.001), regardless of serotonin stimulation, and males (Figure 7D2, tone recall, virus, F_(1,14)_ = 35.20, p<0.0001; Figure 7D3, context recall, t_(14)_ = 4.14, p<0.01). Again, we found that in the absence of chemogenetic inhibition, previous stimulation of serotonin in the adBNST enhanced tone retrieval in females (Uncorrected Fisher’s LSD: ChR2+EYFP vs. ChR2-EYFP, tone-bin 1, p<0.05; Tukey’s multiple comparison test, average tones 1-3, p<0.05). To verify that the DREADD group could acquire fear conditioning without CNO-induced inhibition, ChR2-hM4Di mice were reconditioned following an infusion of aCSF into the adBNST (Figure S6A). Both sexes exhibited higher baseline freezing during the second conditioning session than the first (Figure S6C1, females, t_(4)_ = 3.59, p<0.05; Figure S6D1, males, t_(6)_ = 2.81, p<0.05), indicating that some contextual fear memory was acquired during the first training session. Females also froze more to tone 1 during reconditioning than initial conditioning (Figure S6C1, CNO x tone, F_(4,36)_ = 8.97, p<0.0001), which was not found in males (Figure S6D1, CNO x tone, F_(4,47)_ = 1.75, p = 0.15). Importantly, both sexes exhibited stronger recall of the tone (Figure S6C2, females, CNO, F_(1,10)_ = 18.43, p<0.01; Figure S6D2, males, CNO, F_(1,12)_ = 23.56, p<0.001) and context (Figure S6C3, t_(5)_ = 3.672, p<0.05; Figure S6D3, t_(6)_ = 4.13, p<0.001) after reconditioning, indicating that the underlying circuits remain intact. Together, these results indicate that the CeA-to-BNST pathway plays a necessary role in auditory and contextual fear conditioning in both sexes independently of serotonin.

**Figure 7.**
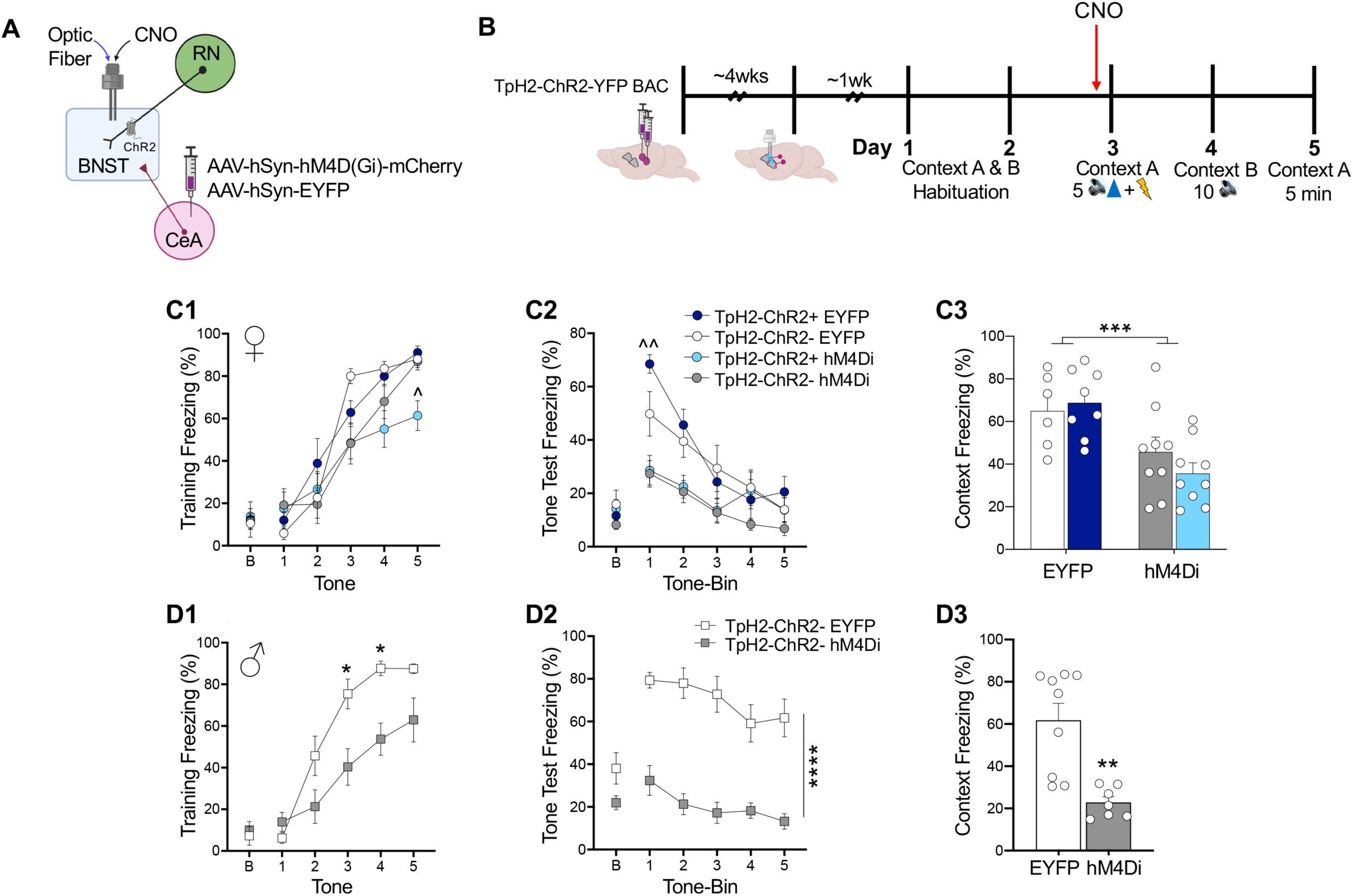
Inhibition of projections from the CeA to the adBNST impairs auditory and contextual fear conditioning in both sexes. **(A-B)** Schematic of procedures. Mice of both sexes were injected bilaterally the CeA and cannulas were implanted bilaterally above the adBNST. CNO was infused directly into the adBNST immediately before auditory fear conditioning, during which blue light was delivered to the adBNST with each tone presentation. Mice were tested to the tone and training context on the following two days without CNO, laser stimulation, or shock. **(C-D)** Chemogenetic inhibition of CeA input to the adBNST impaired acquisition of auditory fear conditioning in **(C1)** females with concurrent stimulation of serotonin in the adBNST and **(D1)** non-stimulated males. During testing, inhibited mice of both sexes exhibited impaired recall of **(C2, D2)** tone and **(C3, D3)** contextual fear memory. Females: n = 6 ChR2-EYFP; n = 8 ChR2+EYFP; n = 9 ChR2-hM4Di; n = 9 ChR2+hM4Di; Males: n = 9 ChR2-EYFP; n = 7 ChR2-hM4Di. Data are represented as mean ± SEM. * p<0.05, **p<0.01, ****p<0.0001, ^p<0.01 vs. all other groups, ^^p<0.0001 vs. ChR2+hM4Di.

### Increasing 5-HT elevates adBNST-to-CeA high gamma communication in females during retrieval

We next used awake behaving electrophysiology to investigate whether increasing serotonin neurotransmission in the adBNST during learning leads to a sex difference in adBNST-CeA communication. Serotonin was stimulated in the adBNST during auditory fear conditioning in male and female mice of both genotypes, as described above (Figure 3A), and local field potential (LFP) signals were recorded in the adBNST and CeA the next day during the tone test (Figure 8A-8C). We focused our analysis on the high gamma frequency range based on evidence indicating that high frequency oscillations (110-160Hz) in this circuit are prominent [49] and gamma oscillations in the extended amygdala contribute to emotion regulation [50]. High gamma was isolated by filtering data between 70-150Hz and power was calculated for the 90-140Hz band (Figure 8D). In females, tone-evoked high gamma power in the adBNST significantly decreased from pre-tone levels in ChR2-mice (Wilcoxon t-test, p=0.012), but not ChR2+ mice (Figure S7E; Wilcoxon t-test, p=0.426). This resulted in a trend towards more tone-evoked high gamma power in the BNST of ChR2+ than ChR2-females (Figure 8E1-2, Mann-Whitney, p=0.077). Notably, these changes were specific to the high gamma band, as there were no differences between genotypes in tone-evoked low gamma (filtered 20-55Hz, calculated for 30-50Hz, Figure 8D) in the adBNST (Figure S7A2, Mann-Whitney, p=0.37), which significantly decreased from pre-tone levels in both groups of females (Figure S7F). There was also more tone-evoked high gamma power in the CeA of ChR2+ than ChR2-females (Figure 8E1-2, Mann Whitney, p=0.051), which again was not found with low gamma (Figure S7A1, Mann-Whitney, p=0.53). In males, tone-evoked high and low gamma in the adBNST significantly decreased from pre-tone levels in both genotypes (Figure S7E-F). In contrast to females, there were no differences between genotypes in tone-evoked high gamma (Figure 8F1-2) or low gamma power (Figure S7C) in either the adBNST (high gamma, Mann Whitney, p=0.61; low gamma, p = 0.41) or the CeA (high gamma, Mann Whitney, p>0.999; low gamma, p = 0.91). Given that GABAergic cells are key for generating activity in the gamma frequency range [51,52], these results suggest that increases in serotonin in the adBNST during fear learning leads to increased GABAergic activity in the adBNST and CeA during fear memory retrieval in females only.

**Figure 8.**
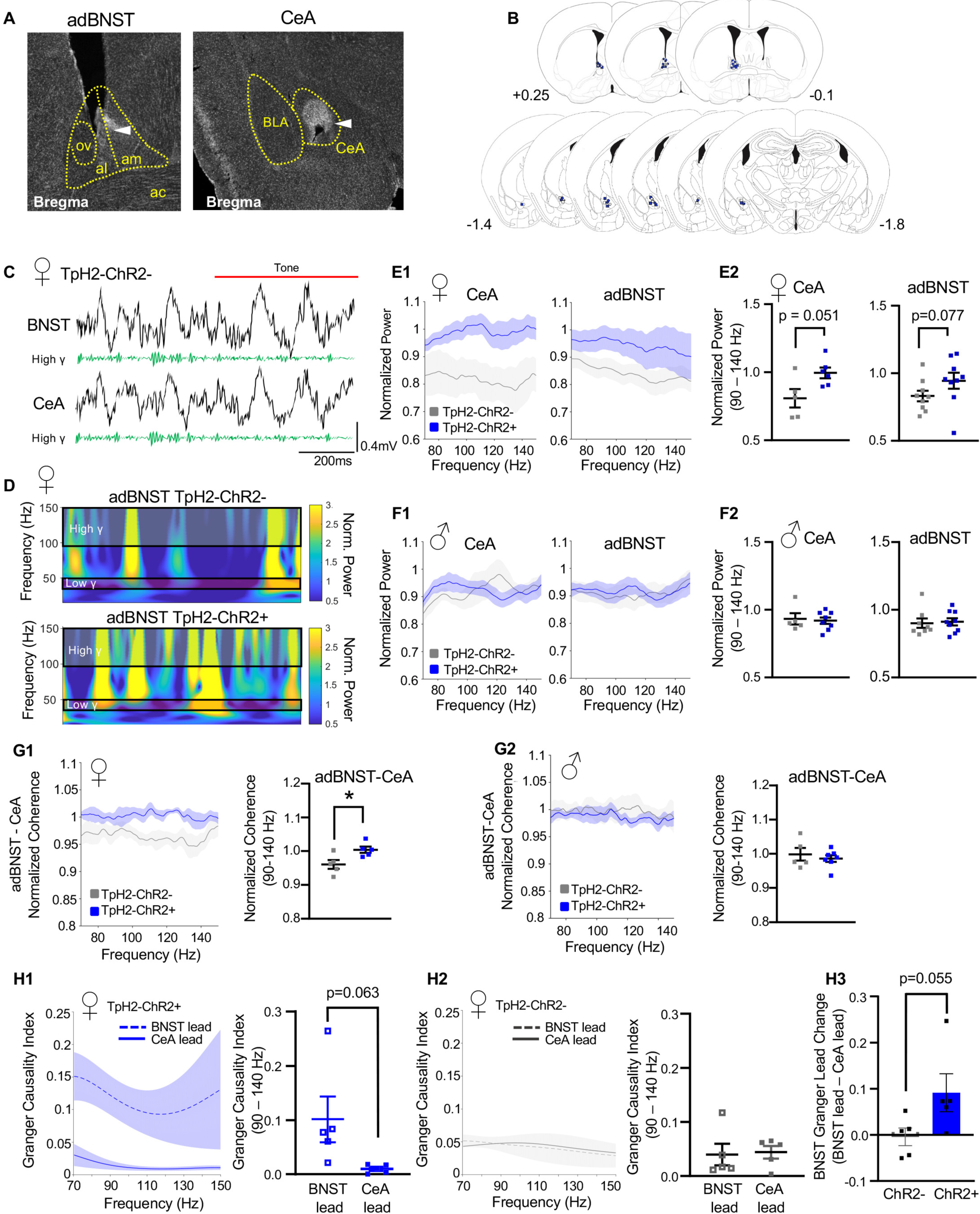
Serotonin increases high gamma synchrony in the adBNST-CeA circuit of female mice during retrieval. **(A)** Example placements of LFP electrodes in the BNST (left) and CeA (right). White arrows indicate electrode tip. **(B)** Mapping of electrode placements in the BNST (top) and CeA (bottom) of all animals. **(C)** Example of a raw LFP (black) and a high gamma filtered LFP recording (green, 70-150Hz) from the BNST and CeA of a female ChR2-mouse. The red line denotes the start of the tone. **(D)** Spectrograms of averaged tone-evoked responses in the BNST of ChR2-(top) and ChR2+ (bottom) female mice. The first four trials of fear memory retrieval were normalized to the average pre-tone periods. The high (90-140Hz) and low (30-50 Hz) gamma bands used in subsequent calculations are highlighted. **(E1-2)** Tone-evoked CeA and adBNST power spectra showing high gamma in ChR2-(grey) and ChR2+ (blue) female mice. In the CeA, there was more normalized high gamma power in ChR2+ females (n=6) than ChR2-females (n=5). A similar effect was found in the BNST (n=9/genotype). **(F1-2)** Tone-evoked CeA and adBNST power spectra showing no difference between ChR2-(grey) and ChR2+ (blue) male mice in normalized high gamma power in the CeA (n = 8 ChR2+; n = 5 ChR2-) or the adBNST (n = 9 ChR2+; n = 8 ChR2-). **(G1-2)** Tone-evoked adBNST-CeA high gamma coherence spectra in **(G1)** female **(G2)** and male mice. Tone-evoked high gamma coherence was significantly higher in ChR2+ (blue) females (n = 5) than ChR2-(grey) females (n = 5), whereas there was no difference between genotypes in males (n = 7 ChR2+; n = 5 ChR2). **(H1-H3)** Tone-evoked Granger Causality index for high gamma adBNST lead (broken line) and CeA lead (solid line) in **(H1)** ChR2+ and **(H2)** ChR2-female mice (n=5/genotype). In ChR2+ females, the probability of a BNST lead in high gamma was higher than a CeA lead, whereas neither site led the other in ChR2-females. **(H3)** The difference between adBNST and CeA lead shows a significant increase in the strength of the adBNST lead in ChR2+ than ChR2-female mice. SEM is shown in shaded regions. *p < 0.05.

Next, we used coherence to investigate high gamma synchrony in the adBNST-CeA circuit. This analysis showed a significant elevation in tone-evoked adBNST-CeA high gamma coherence in ChR2+ (n=5) relative to ChR2-(n=5) female mice (Figure 8G1, Mann Whitney, p=0.032). Notably, this increase in tone-evoked coherence was also specific to the high gamma range, as no changes in low gamma (30-50 Hz) BNST-CeA coherence were observed (Figure S7B, Mann Whitney, p=0.931). Furthermore, in male mice there were no differences between genotypes in adBNST-CeA high or low gamma coherence (Figure 8G2, high gamma, ChR2+ (n=7) vs ChR2-(n=5), Mann Whitney, p=0.999; Figure S7D, low gamma, Mann Whitney, p=0.527).

Having established a serotonin-induced change in high gamma BNST-CeA coherence in females, we next used Granger causality [53] to assess directionality of high gamma communication in this circuit. This analysis showed that in ChR2+ females, the probability of a BNST lead was higher than a CeA lead in gamma frequency (Figure 8H1-2, Mann Whitney, p=0.063), whereas neither site led the other in ChR2-females. Consequently, there was an increase in the strength of the adBNST lead in ChR2+ compared to ChR2-females (Figure 8H3). Thus, in females, but not males, increasing serotonin in the adBNST during fear learning leads to greater tone-evoked high gamma adBNST-CeA synchrony and increased adBNST-to-CeA high gamma communication during fear memory retrieval.

## DISCUSSION

Serotonin is implicated in PTSD [54,55] and modulates fear learning [56–58], but it is not known if it contributes to sex differences in PTSD risk [3,59,60]. Using auditory fear conditioning as a model of fear learning in mice, we first identified sex differences in response to brain-wide elevations in serotonin, with female mice exhibiting higher sensitivity to SSRI-induced increases in tone recall than males. Our finding that a lower SSRI dose enhanced training-induced upregulation of c-Fos expression in the adBNST and CeA of females, but not males, implicated the extended amygdala in this sex difference. Selectively increasing serotonin in the adBNST during conditioning with optogenetics led to similar changes in c-Fos expression in the extended amygdala of females and was sufficient to increase tone recall in females only, behavioral effects that we demonstrate were mediated by activation of 5-HT_2C_ receptors in the adBNST. In line with these findings, our analysis of different serotonin receptors in the adBNST revealed a sex difference in 5-HT_2C_ receptors only, with higher transcript levels found in females than males. Electrophysiological analyses also revealed a sex difference, whereby stimulating serotonin terminals in the adBNST led to increased adBNST-CeA high gamma (90-140 Hz) synchrony and higher probability of an adBNST-to-CeA communication pattern in this frequency range in females only. Collectively, these results provide the first evidence of a sex difference in fear learning mediated by the raphe-adBNST-CeA circuit. In humans, such sex differences may contribute to the increased risk of PTSD found in women compared to men.

Our citalopram findings are in agreement with previous studies conducted by our group and others demonstrating that acute SSRI treatment prior to auditory fear conditioning increases tone recall [13–15] and immediate early gene expression in the BNST and CeA of male rats [14,15] . Here, using mice we show that females are more sensitive to these effects than males, as indicated by our finding that a lower dose of citalopram (10mg/kg) was sufficient to induce these changes in females but not males. Interestingly, this same dose of citalopram that was not effective in male mice was effective in male rats, which like our study, was administered 60 minutes before conditioning [13]. We attribute this discrepancy to a species difference in drug metabolism, which is faster in mice than rats [61]. There are also species differences in drug metabolizing enzymes [62], including those involved in SSRI metabolism [63]. It is therefore possible that 10mg/kg of citalopram had a weaker effect on serotonin levels in male mice than male rats, which is why a higher dose (20mg/kg) was required to induce behavioral effects similar to those described in rat. The advantage of using an SSRI dose that was without an effect in male mice is that it allowed us to detect a sex difference in serotonin sensitivity. To our knowledge, this is the first SSRI study to test acquisition of fear conditioning in female rodents.

Consistent with our SSRI findings, we found a sex difference in the effects of selectively activating serotonin inputs to the adBNST, which enhanced tone recall, but not context recall, in females only. In contrast, Marcinkiewcz et al. reported that optogenetic stimulation of serotonin terminals in the BNST increases recall of both tone and context in males. Although the stimulation protocol (10 mW, 473 nm, 5 ms pulses, 20 Hz) was the same in both studies, other methodological differences may account for our different results. In the Marcinkiewcz et al. study, the optic fiber positioning increased the likelihood that they also affected serotonin in the ventral BNST. Given that the ventral BNST is implicated in recall of contextual fear conditioning via projections to the ventral tegmental area [64], it is feasible that the additional involvement of serotonin in this region affected fear learning in males. Unlike our study, Marcinkiewcz et al. did not habituate mice to training or testing contexts prior to fear conditioning, both of which were consequently novel to their animals during fear conditioning and tone recall. Novelty increases corticosterone [65] and activity in the BNST [66,67] and BLA [68], which may have led to activation of BNST circuits that differed from our study. It is also noteworthy that Marcinkiewcz et al. used a viral strategy to express ChR2 in serotonergic neurons of Sert-Cre mice and we used a BAC transgenic mouse line that expresses ChR2-EYFP in serotonin neurons.

Serotonin has different effects on cell excitability in the BNST that are dependent on which receptor has been activated [69]. While activation of 5-HT_1A_ receptors in the BNST mediates an inhibitory response, activation of 5-HT_2A_, 5-HT_2C_, and 5-HT_7_ receptors mediate excitatory responses [25,69,70]. Consequently, the effects of serotonin on anxiety vary, with activation of 5-HT_1A_ receptors in the BNST being anxiolytic [46] and activation of 5-HT_2C_ receptors being anxiogenic [25]. The 5-HT_2C_ receptor also plays a role in fear learning, as demonstrated by the finding that systemic administration [71] or direct infusions of a 5-HT_2C_ antagonist into the BNST [14] block SSRI-induced increases in tone recall in male rats. Here, we extend these findings to females and show that blocking these receptors blocks the effects of optogenetically increasing serotonin in the adBNST on tone recall. In addition, we found that serotonin receptor transcript levels in the adBNST were the highest for the 5-HT_1A_ and 5-HT_2C_ receptors in both sexes, but that the expression of 5-HT_2C_ receptors and the relative expression of 5-HT_2C_ to 5-HT_1A_ receptors was higher in females than males. As a result, it may be more likely that serotonin release activates 5-HT_2C_ receptors in females than males, in turn increasing risk of anxiogenic effects and enhanced fear memory in females. In contrast to our study, which is the first to evaluate sex differences in the expression of numerous serotonin receptors in the adBNST, recent work focusing on 5-HT_2C_ receptors reported no sex difference in BNST *Htr2c* transcript levels [72]. Unlike our study, which used naïve animals, Flanigan et al. measured *Htr2c* transcript following 3-weeks of binge alcohol consumption. Given that alcohol consumption affects 5-HT_2C_ receptor activity [73], behavioral testing may have affected 5-HT_2C_ expression, potentially obscuring detection of sex differences. Their inclusion of the ventral BNST, a region excluded in our analysis, may also have contributed to the discrepancy in our findings, particularly if the sex difference we report is specific to the adBNST.

In males, we consistently found that increasing serotonin (locally or systemically) during conditioning reduced pre-tone freezing the next day, indicating that serotonin reduced generalization of fear to the recall context. In contrast, there was less fear generalization in females, whose pre-tone freezing was low regardless of serotonin manipulation. Considering that we used two separate environments for training and tone recall that differed in shape (square versus rectangle), floor texture, and smell, freezing to the context used for tone recall was unexpected. Our findings reveal both a sex difference in fear generalization and a role for serotonin in the adBNST in reducing this response. BNST lesions have been shown to affect generalization of cued fear [74], but less is known regarding the role of this region in generalization of contextual fear.

Auditory fear conditioning studies typically do not focus on the BNST, which can be lesioned or functionally inactivated without affecting acquisition [14] or recall [75] of cued fear memory. Instead, the BNST is more widely known for its role in anxiety-like behavior [37,76]. However, accumulating evidence supports a role for the BNST in cued fear learning. For example, chemogenetic activation of GABAergic or somatostatin neurons in the BNST during either auditory fear conditioning or fear memory consolidation enhances tone recall [33]. It has also been shown that auditory fear conditioning upregulates c-Fos expression in the BNST of male mice [33], effects that we replicated in both sexes and found most consistently in the anteromedial subregion. Single cell recordings reveal that neurons in the adBNST become responsive to the conditioned stimulus after auditory fear conditioning, with different subpopulations of cells developing excitatory or inhibitory responses [77]. Here, we report that the conditioned stimulus evoked a decrease in high and low gamma in the adBNST of both sexes during recall. Interestingly, stimulation of serotonin terminals in the adBNST during conditioning blocked this decrease in females but not males. Importantly, this effect cannot be attributed to a sex difference in the number of serotonin projecting neurons to the adBNST, which we show was similar in males and females in both the dorsal and median raphe nuclei.

The adBNST and CeA are reciprocally interconnected brain regions, both of which are key components of the extended amygdala. GABAergic projections originating primarily from the alBNST and ovBNST [78] terminate in the medial subdivision of the CeA (CeM), with sparse projections terminating in the lateral subdivision (CeL) [43] . Conversely, the CeL sends dense GABAergic projections to the adBNST, many of which terminate in the ovBNST [79]. Despite substantial evidence indicating the involvement of both regions in fear learning, few fear conditioning studies have investigated adBNST-CeA communication *in vivo*, and none characterize modulation by serotonin or include females. We focused our analysis on gamma oscillations, because they are mediated by GABAergic neurons, which largely comprise both structures and are implicated in long-range synchrony during cognitive tasks [51,80,81]. Consistent with previous reports demonstrating high frequency oscillations (110 – 160Hz) in the adBNST and CeA of the rat [49], we saw task-evoked changes in the high gamma band (90-140Hz) in this circuit. We show that in females only, stimulation of serotonin in the adBNST during fear learning increased adBNST-CeA high gamma coherence and increased the probability that information was transferred in an adBNST-to-CeA direction during recall. The increase in tone-evoked high gamma oscillations in stimulated females suggests that there was serotonin-mediated plasticity in a subset of GABAergic adBNST neurons that impacted communication with the CeA. We previously showed that activation of somatostatin (SST) interneurons in the BLA during recall of a non-threatening stimulus increases BLA high gamma power, leading to suppressed fear expression [82,83] . Conversely, activation of SST neurons in the adBNST drives consolidation of fear memory [33], suggesting that the fast gamma we observe in females during elevated fear recall may be related to serotonergic activation of SST neurons. Given that high gamma oscillation patterns reflect local interneuron activity [51], a better understanding of connectivity patterns between locally- and distally-projecting GABAergic adBNST neurons is necessary for understanding how increased high gamma adBNST-to-CeA communication drives fear memory. One possibility is that a serotonin-induced increase in GABAergic activity in a subset of locally-projecting adBNST neurons rhythmically inhibits CeA-projecting GABAergic adBNST neurons, thereby disinhibiting the CeM - the main target of adBNST projections [43] and a key region for fear expression [84]. Notably, 5-HT_2C_ receptors in the adBNST are located on locally-projecting interneurons [25], making them likely candidates for mediating such CeM disinhibition.

Collectively, our data indicate that females are more sensitive than males to the enhancing effects of serotonin on fear memory because they have higher activation of 5-HT_2C_ receptors in the adBNST during learning, which leads to stronger adBNST-to-CeA high gamma communication during recall. Additionally, our finding that fear learning is dramatically impaired by blocking projections from the CeA to the adBNST highlights the importance of communication between these regions for fear learning in both sexes, even in the absence of serotonin manipulation. The sex difference we describe in the raphe-BNST-CeA circuit and consequent effects on fear learning may account for the increased risk of PTSD found in women compared to men. This work highlights the importance of including both sexes in preclinical research, as it may better inform neural mechanisms underlying sex differences in clinical diagnoses.

## Acknowledgements

We thank members of the animal care staff at Hunter College, particularly Barbara Wolin and Sonia Acevedo. We thank Sadiyah Hanif for assistance with tissue processing. We would also like to acknowledge Dr. Glenn Schafe and Dr. Alvaro Garcia-Garcia for their helpful comments and suggestions. This project was supported by NIMH R21MH114182 (N.S.B. & E.L.), National Institute on Minority Health and Health Disparities G12MD007599 (N.S.B.), NIMH R01MH118441 (E.L.) & PSC-CUNY Awards (N.S.B. & E.L.).

## Author Contributions

R.R. performed the SSRI, 5-HT_2C_ antagonist and chemogenetic experiments. R.R. and Jinah.L. performed the optogenetic and *in vivo* electrophysiology experiments. Jia L. performed the qRT-PCR experiments. R.R., C.F.H., E.L. and N.S.B analyzed the data. R.R. and N.S.B. designed the experiments. E.L. and N.S.B. supervised the study. R.R., E.L. and N.S.B wrote the manuscript with help from other authors.

## Methods

### Animals

Male and female SJL-Tg (Tph2-COP4*H134R/EYFP) 5Gfng/J mice (C57BL/6J background) (Jackson Laboratory, Bar Harbor, ME) were bred in-house (Hunter College, New York, NY) and used in all experiments. In this mouse line, channelrhodopsin (ChR2) – enhanced yellow fluorescent protein (EYFP) is directed to 5-HT neurons by the TpH2 promotor of the BAC transgene. In the dorsal raphe nucleus (DRN) and median raphe nucleus (MRN), ChR2-EYFP+ neurons are exclusively TPH2+, confirming that ChR2-EYFP is selectively expressed in serotonergic neurons. TPH2+ immunofluorescence also strongly colocalizes with EYFP+ neurons in the DRN and MRN (81-85%) [85]. Upon weaning, ChR2-EYFP mice (ChR2+) and littermate controls (ChR2-) of the same sex were group housed (2 – 4/cage) with food and water available *ad libitum* and maintained on a 12-hour light/dark cycle. All procedures were conducted during the light cycle and approved by the Hunter College Institutional Animal Care and Use Committee.

### Drug Administration

Citalopram hydrobromide (Sigma-Aldrich, #5062380001, St. Louis, MO) was dissolved in 0.9% sterile saline and injected intraperitoneally (i.p.) at a dose of 20 mg/kg or 10 mg/kg [13,14] either 60 minutes before fear conditioning or 160 minutes before transcardial perfusion. The selective 5-HT_2C_ receptor antagonist RS102221 (Tocris Bioscience, #1050, Minneapolis, MN) was dissolved in a 50% aCSF and 50% sterile saline solution and infused bilaterally into the aBNST at a dose of 0.5 µg per side [14][14]. CNO (Millipore Sigma, #C0832, St. Louis, MO) dissolved in artificial cerebrospinal fluid (aCSF) (Tocris Bioscience, #3525, Minneapolis, MN) was infused bilaterally into the adBNST at a dose of 1mM per side. For local drug infusions a volume of 0.25µL was infused bilaterally at a rate of 0.1µL/min into the adBNST (GenieTouch, Kent Scientific, CT) 10-15 minutes prior to fear conditioning. Injection cannulas (P1 Technologies, Roanoke, VA) were left in place for an additional 5 minutes after the infusion to ensure diffusion of drug away from the cannula tip.

### Estrous Cycle Monitoring

Vaginal cytology was used to monitor estrous cycle, as previously described [86]. Cells were collected with a cotton swab soaked in 0.9% sterile saline that was shallowly inserted into the vaginal canal. The swab was rolled onto a blank slide and cells were viewed with a bright field microscope. Cell composition was used to determine phase of estrous cycle in the following way: keratinized needle-like, nucleated epithelial cells and leukocytes indicated metestrus; high levels of leukocytes with some cornified cells indicated diestrus; small rounded cells that were often clustered indicated proestrus; and cells that were primarily cornified indicated estrus . A female was classified as “High Estradiol” if cytology indicated proestrus or estrus and “Low Estradiol” if cytology indicated metestrus or diestrus. Each day, swabbing was conducted in females housed in their home cage a minimum of 60 minutes before being placed in Context A or B.

### Stereotaxic Surgery

Nine to ten-week-old male and female ChR2+ and ChR2-mice were anesthetized with 2% isoflurane in oxygen, placed in a stereotaxic frame (Kopf Instruments, Tujunga, CA) and maintained on 1.5% isoflurane throughout surgery. Temperature was maintained at 37°C ± 1°C with a feedback regulated heating pad. Animals received dexamethasone (0.05mL, 1mg/mL, s.c.) and bupivacaine under the scalp (0.1mL, 5mg/mL, s.c.) and craniotomies were made using anteroposterior (AP) coordinates from bregma, mediolateral (ML) coordinates from the midline, and dorsoventral (DV) coordinates from the brain surface. For optogenetic experiments, fiber optics were bilaterally implanted in the adBNST (+0.25 – +0.35 mm AP, ± 1.00 mm ML, - 4.15 mm DV) of mice. Fiber optics were constructed with ceramic ferrules (ThorLabs, CFLC230-10, Newton, NJ) or ordered precut (Hangzhou Newdoon Technology Co., FOC-C-1.25-200-6.0-0.22, China) and secured with opaque C&B Metabond (Parkell, Edgewood, NY) and dental cement (Teets, Co-Oral-Ite Dental Mfg, Diamond Springs, CA). For electrophysiological recordings, tungsten wires (California Fine Wire, #100211) were placed bilaterally into the CeA (−1.17 mm AP, ± 2.4 mm ML, -4.4 mm DV), super glued to fiber optics or cannulas and placed bilaterally into the adBNST (+0.25 – +0.35 mm AP, ± 1.00 mm ML, - 4.15 mm DV), and secured with Metabond and black dental cement. Skull screws secured over the frontal cortex served as reference and screws over the cerebellum served as ground. Wires were connected to a 16-channel custom built electronic interface board carrying an Omnetics connector (Neuralynx, A79018-001, Bozeman, MT). Tungsten wires were gold-pinned (large, Neuralynx, Bozeman, MT) to the board and reference and ground wires were soldered to their respective locations on the board. For DREADD experiments 6-7 week old mice were injected bilaterally with the inhibitory virus AAV-hSyn-hM4D(Gi)-mCherry (AddGene, 50475-AAV2) or control virus AAV-hSyn-EYFP (AddGene, 50465-AAV2, Watertown MA) into the CeA (−1.17 mm AP, ± 2.4 mm ML, -4.4 mm DV) at a rate of 0.1µL/min with 10µL Hamilton syringes (QSI Stereotax Injector, Stoelting, IL). Following injection (0.25µL/side), the skull was closed with bone wax and the incision was sutured. Approximately four weeks later, mice were implanted bilaterally with custom cut 26-gauge guide cannulas (P1 Technologies, VA) above the adBNST (+0.25 - +0.35 mm AP, ±1.00 mm ML, -3.75 mm DV) secured with opaque C&B Metabond (Parkell, Edgewood NY) and black dental cement (Teets, Co-Oral-Ite Dental Mfg, Diamond Springs CA). Dummy cannulas were inserted prior to surgical recovery. For the 5-HT_2C_ receptor antagonist experiments, there were no viral injections but the same procedures were used for implanting cannulas. Postoperatively, all animals received 0.25mL sterile saline (s.c.) and 0.08mL carprofen for pain relief (1mg/mL, i.p.). Mice were group housed in cages warmed by a heating pad until recovery. Placements of fiber optics, cannulas and wires were confirmed upon completion of experiments. For tungsten wires, this required creating a lesion by passing current (0.05mA, 25s) through the channels of anesthetized animals.

### Fear Conditioning

Adult (∼12 weeks old) mice were fear conditioned in plexiglass chambers with aluminum walls and a stainless steel rod floor capable of delivering scrambled foot shock (Context A) (Med Associates, St. Albans, VT). Overhead lamps maintained light levels at ∼30 lux and the conditioning box was cleaned with ethanol between animals. During the cued recall test, mice were placed in a custom constructed gray wood box with high walls and a smooth floor (∼70 lux) (Context B). Orange scented Clorox wipes were used to clean the recall box between animals. Behavior and electrophysiological data were recorded using an infrared OptiTrack camera and Neuromotive software running in conjunction with Central software (Blackrock Microsystems, Salt Lake City, UT). Timestamped data were extracted from Central-generated NEV files in MATLAB and analyzed off-line. For DREADD and antagonist experiments animals were handled for 5 days (5 min/day) and acclimatized to injection system prior to fear conditioning. For all other experiments following 3 days of handling (5 minutes/day), mice were habituated for 20 minutes to the training (Context A, day 1) and testing context (Context B, day 2). On day 3, mice were fear conditioned in Context A with 5 presentations of a tone conditioned stimulus (CS: 2 kHZ tone, 85 dB, 30 sec) that co-terminated with a footshock unconditioned stimulus (US: 0.7 mA, 2sec) with an inter-trial interval that varied randomly between 90 – 120 seconds. The next day (day 4), cued recall was tested in Context B with 10 presentations of the tone in the absence of shock. The properties of the tones and inter-trial interval were identical during conditioning and testing. On day 5, animals were returned to the training context (Context A) for 5 minutes and contextual fear memory was evaluated. Time spent freezing to tone and context (immobility unrelated to respiration) was hand-scored by an experimenter blind to group. Freezing during the 30-sec period before the first tone test was considered baseline freezing. Freezing behavior is reported as a percentage of total tone time, pre-tone time, and time in the training context.

### Optogenetic Stimulation

Optogenetic stimulation of 5-HT terminals in the adBNST involved delivery of ∼10mW of blue light (473 nm, 5 msec pulses, 20 Hz) (Laserglow, Toronto, Canada; WaveForm 33500B, Keysight Tech, CO). This stimulation protocol has previously been shown to elicit optimal responses in 5-HT neurons [85,87] and effectively modulate behavior [46] in this mouse line. During fear conditioning, blue light was delivered only during each tone presentation (5 tones, 30 seconds). In naïve (non-trained) animals, blue light was delivered 5 times for 30 seconds at intervals that varied pseudorandomly between of 30 – 60 sec.

### Retrograde Tracing

A 10µL Hamilton syringe (QSI Stereotax Injector) containing 0.25 µL of the retrograde tracer Cholera Toxin Subunit B – Alexa Fluor 647 (CTB) (Thermofisher Scientific, #C34778) was fitted with a fine tipped glass capillary tube and lowered into the right adBNST (+0.25 –+ 0.35 mm AP, - 1.00 mm ML, - 4.15 mm DV). CTB was infused at a rate of 0.1µL/min (Stoelting) and left in place for 7 minutes. The craniotomy was closed with bone wax (Look, Surgical Specialties) and skin was closed with nonabsorbable nylon suture (5-0, Butler Shein). Ten days later, animals were perfused and every 6^th^ section containing the extended amygdala and raphe nuclei was washed in PBS-0.1% triton (PBST) and incubated in 10% normal donkey serum in PBST for 1 hour at RT. Sections were incubated in rabbit anti-EYFP (1:500, ThermoFisher Scientific, #A11122), mouse anti-GAD65/67 (1:250, Santa Cruz Biotechnology, #sc-365180), and guinea pig anti-vGlut3 (1:250, Synaptic Systems, #135204) in 1% normal donkey serum in PBST for 48 hours at 4°C. Then slices were washed in PBS and incubated in donkey anti-rabbit 488 (1:100, Jackson ImmunoResearch, #711545152), donkey anti-guinea pig Dylight 405 (1:100, Jackson ImmunoResearch, #706475148), and donkey anti-mouse 594 (1:200, Jackson ImmunoResearch, #715585150) for 1 hour at RT. Mounted sections were cover slipped with ProLong Gold antifade mounting medium (ThermoFisher Scientific, #P36930). Sections were imaged using a Nikon A1 confocal microscope (Melville, NY). Cell counting was done manually using ImageJ.

### C-Fos Immunohistochemistry

Ninety minutes after conditioning, mice were deeply anesthetized with a mixture of ketamine (100 mg/kg, i.p.) and xylazine (7 mg/kg, i.p.) and transcardially perfused with cold PBS followed by 4% paraformaldehyde in PBS. Brains were removed, postfixed in 4% paraformaldehyde overnight, and then cryoprotected in 30% sucrose for at least 5 days. A cryostat (CM3050S, Leica, Germany) was used to cut 35 µm coronal sections. Every 6^th^ section containing the extended amygdala was processed for c-Fos immunoreactivity. Free-floating sections were washed in PBS (3x, 10min) followed by PBS-1.0% triton (PBST, 30 min) and then blocked in 5% normal donkey serum (NDS) in PBST for 1hr at room temperature. Tissue was incubated overnight in anti-c-Fos antibody (rabbit polyclonal, 1:1k, Abcam, #190289, Cambridge, MA) in 5% NDS in PBST at 4°C. Sections were then washed in PBS and incubated in donkey anti-rabbit Alexa Fluor 488 secondary antibody (1:500, Thermo Fisher Scientific, #A11055, Waltham, MA) in PBS for 1hr at room temperature. Mounted sections were cover slipped with ProLong Gold plus DAPI antifade mounting medium (ThermoFisher Scientific, #P36931, Waltham, MA) and imaged with a Nikon A1 confocal microscope (Melville, NY). Cell counting was done manually using ImageJ.

### GFP Immunohistochemistry

Following transcardial perfusion brains were removed and post fixed as previously described. A cryostat was used to cut 50 µm coronal sections. Every 6^th^ section containing the extended amygdala and raphe nuclei was washed in PBST then blocked in 10% NDS-PBST for 1 hour at RT. Sections were incubated in rabbit anti-EYFP (1:500, ThermoFisher Scientific, #A11122) in 1% NDS-PBST at 4°C for 48 hours. Sections were washed in PBS and incubated in donkey anti-rabbit 488 (1:100, Jackson Immunoresearch, #711545152) for 1 hour at RT. Mounted sections were cover slipped with ProLong Gold antifade and imaged with a Nikon A1 confocal.

### RNA Extraction, cDNA Synthesis and qRT-PCR

Bilateral tissue punches (1 mm) from the adBNST of 12 – 13 week-old male and female ChR2-mice were flash frozen for subsequent processing at the Epigenetics Core of CUNY Advanced Science Research Center. Briefly, RNA was extracted with Trizol (Thermofisher Scientific, #15596026) and purified using the Qiagen RNeasy Micro kit (Qiagen, #74181) following the manufacturer’s protocol. RNA was reverse transcribed with qScript cDNA supermix (Quanta Bio, #95048-500) and qRT-PCR was performed with QuantaStudio 7 Real-time qPCR System (Thermofisher, #A43183). Transcript levels of *Htr2c, Htr1a, Htr7, Htr2a, Htr3a,* and *Sst* were quantified in triplicate. The average values for each transcript was calculated after normalization to *Gapdh* and *18s* using primers listed in Reagent Table.

### Awake-Behaving Electrophysiology

Data were analyzed only from channels with confirmed BNST and CeA placements (Figure 7A-B). During behavior, local field potentials were sampled from both sites at 2kHz with a low pass filter (1kHz). Data were imported into Matlab (Natick, MA) and custom-written scripts were used for analyses. Line noise (60Hz) and its harmonics were filtered from the signal (*removeLineNoise_SpectrumEstimation,* [88]. Time stamps of interest (30 sec of each tone and 30 sec of each accompanying pre-tone) were extracted. Pre-tone was used to normalize electrodes across both sites and animals for tone-evoked power and coherence. For females, pre-tone activity was an average of 30-sec prior to all 10 tones. For males, to account for the high contextual fear seen early in the recall session (Figure S4), pre-tone activity was an average of 30-sec prior to tones 3-10. The Morlet wavelet transform (q=3, for frequencies between 15 and 150 Hz, Figure 8D) was used to assess whether a high-vs low-frequency gamma band changed with behavior [49]. A custom-written script (mtcsg, provided by K. Harris and G. Buzsaki) was used to calculate multi-taper spectral power with a 250 ms (500 samples) moving window, 240 ms (480 samples) overlap, a time-bandwidth product of 1.5 with 2 tapers, and 2048 FFT in BNST and CeA during pre-tone and tone presentations, allowing for resolution of about 1Hz. The 30-sec tone period was cut into 1-sec periods, and power was calculated for each second, averaged across the 30 sec, and normalized to the average of the pre-tone periods. If the power of the estimated signal exceeded the SD of the median by 15x in a given 1-sec period, that second was counted as noisy and eliminated from the analysis. Coherence between ipsilateral (left) adBNST and CeA was determined using multitaper coherence (mtchg, provided by K. Harris) and the same windowing parameters as for the power analysis listed above. The CS-evoked signal was normalized to pre-tone data, as described above. Granger causality analysis was performed using the Multivariate Granger Causality (MVGC) Toolbox (v.1.3), as described in Barnett and Seth (2014) [53]. The analysis was run on the first 4 trials, where the behavioral difference was most apparent. The signal was filtered for the 60Hz noise as described above, and each 30-sec trial was analyzed with windows of 100 ms (frequency resolution 0.5Hz), and then the average strength of BNST vs CeA lead across the 4 trials was compared.

## STATISTICAL ANALYSES

Data were analyzed with the non-parametric Mann-Whitney test (electrophysiology), one-way ANOVA, Student’s t-test for independent samples, and two-way repeated measures ANOVA with Tukey’s HSD post hoc tests using GraphPad Prism software (GraphPad, San Diego, CA). A mixed effects model for repeated measures was used to analyze behavioral data with the 5-HT_2C_ antagonist (SPSS v 25.0, IBM, Armonk, NY). Interaction effects are reported if they reached significance. For acquisition data, we only included tone-evoked freezing responses following the first foot shock in the analysis (tones 2-5). For c-Fos data, planned comparisons of trained groups were used to determine whether increasing serotonin augments learning-induced increases in immuno-positive cells. Significance levels were set at p<0.05.

## SUPPLEMENTARY FIGURE LEGENDS

**Figure S1. Stage of estrous cycle does not influence fear conditioning. (A)** Schematic of procedures indicating when vaginal swabs were taken each day. **(B)** Representative images of vaginal cytology of TpH2-ChR2-YFP BAC female mice. **(C)** Estradiol status on the day of auditory fear conditioning did not affect **(C1)** acquisition, **(C2)** tone recall or **(C3)** recall of contextual fear memory (n = 15 low estradiol; n = 12 high estradiol). **(D1)** Estradiol status on the day of the tone test did not affect tone recall (n = 14 low estradiol; n = 13 high estradiol). **(D2)** Estradiol status on the day animals were placed back in the training context (Context A) did not affect recall of contextual fear memory (n = 13 low estradiol; n = 13 high estradiol). Data are represented as mean ± SEM.

**Figure S2. Generalization of fear to the testing context is short lasting in saline-treated males. (A)** In males, acute systemic administration of citalopram prior to auditory fear conditioning reduced pre-tone freezing in a different context the next day (n = 14 per citalopram dose; n = 28 combined saline). The same data are shown as baseline freezing in Figure 1C2. **(B)** Schematic of behavioral procedures. The same two doses of citalopram or saline were injected in a new cohort of male mice before auditory fear conditioning. On the following two days, mice were tested in the testing context (Context B) without exposure to tones and the training context (Context A). **(C1)** Acute citalopram treatment did not affect percentage of time spent freezing to the tone during conditioning (n = 9 saline; n = 7, 10mg/kg; n = 9, 20mg/kg). **(C2)** The next day, drug-free mice were placed in the testing context (Context B) and freezing was measured when tones are usually presented. Saline-treated mice only froze more than citalopram-treated mice at baseline (B) and when the first two tones are usually presented (Time-Bin 1). **(C3)** Citalopram treatment did not affect recall of contextual fear memory. Data are represented as mean ± SEM. ***p<0.001 vs. saline, **p<0.01 vs. saline, *p<0.05 vs. saline.

**Figure S3. 5-HT input to the adBNST is similar in males and females. (A)** Alexa Fluor 647-conjugated cholera toxin subunit B (CTB) was injected into the right adBNST of TpH2-ChR2+ mice and expression was visualized in the raphe nuclei 10 days later. **(B)** Representative images from a female mouse showing **(B2)** TpH2-ChR2-EYFP+ neurons (green) and CTB+ neurons (far red) in the dorsal raphe (DRN) and median raphe nuclei (MRN). **(B1)** High magnification of the DRN showing a co-labeled cell. **(C-F)** Quantification of neurons co-labeled with TpH2-ChR2-EYFP and CTB reveal similarities between males and females in the number **(C, E)** and percentage **(D, F)** of TpH2-ChR2+ neurons in the DRN and MRN that project to the adBNST. n = 7 males; n = 6 females. Data are represented as mean ± SEM.

**Figure S4. Generalization of fear to the testing context is short lasting in non-stimulated males. (A)** In males, optogenetic stimulation of 5-HT terminals in the adBNST during auditory fear conditioning decreased pre-tone freezing in the testing context the next day (n = 22 ChR2+ males; n = 21 ChR2-males). The same data are shown as baseline freezing in Figure 3E2. **(B)** Schematic of behavioral procedures. A separate cohort of TpH2-ChR2-males received light stimulation during auditory fear conditioning but were not exposed to tones in Context B the next day. The following day they were tested in the training context (Context A). **(C1)** During training, freezing to the tone increased with each tone-shock pairing (n = 10). **(C2)** The next day, mice exhibited freezing in the testing context that decreased by the time the tones are usually presented. Baseline freezing in the previous cohort of TpH2-ChR2-(white squares) and TpH2-ChR2+ males (blue squares) (Figure 3E2) is included for purposes of comparison. **(C3).** Freezing to the training context (Context A) is shown for mice that were not exposed to tones on the previous day (ChR2-No Tones) and the previous cohort that was tone tested (ChR2-Tones, ChR2+Tones, Figure 3E3). Data are represented as mean ± SEM. ***p<0.001.

**Figure S5. Effects of the 5-HT_2C_ receptor antagonist on tone recall are temporary. (A)** Schematic of procedures. The 5-HT_2C_ receptor antagonist RS102221 was bilaterally infused into the adBNST of female ChR2+ mice 10-15 minutes before fear conditioning (Train 1). On the following two days, mice were tested to the tone (Tone Test 1) and the training context. Forty-eight hours later, a subset of these animals received an infusion of aCSF into the adBNST before a second fear conditioning session (Train 2) followed by a second tone test (Tone Test 2). Serotonin was stimulated in the adBNST with blue light during both training sessions. **(B-C)** Reconditioning in the absence of the antagonist increased freezing during **(B)** training and **(C)** the tone test. n = 7 RS102221; n = 4 aCSF. Data are represented as mean ± SEM. *p < 0.05, **p < 0.001, ***p< 0.0001.

**Figure S6. CNO-induced impairments in fear conditioning are not permanent. (A)** Schematic of procedures. Following bilateral injection of hM4Di into the CeA, projections from the CeA to the BNST were inhibited in mice of both sexes via direct infusion of CNO into the adBNST before fear conditioning (Train 1). Effects on tone and context recall were tested (Test 1). The same mice received an infusion of aCSF into the adBNST before a second fear conditioning session (Train 2), followed by a second tone and context test (Test 2). **(B)** Representative confocal images showing expression of the hM4Di-DREADD virus (mCherry tagged) (red) and EYFP control virus (green) in the CeA (injection site) (left) and the adBNST (right) in a female mouse. **(C1-D1)** Reconditioning in the absence of inhibition increased (**C2, D2**) tone and (**C3, D3**) context freezing in both sexes. n = 7 males; n = 6 females. Data are represented as mean ± SEM. ov = oval BNST, al = lateral BNST, am = medial BNST, CeA = central nucleus of the amygdala, BLA = basolateral amygdala, ac = anterior commissure. * p < 0.05, **p < 0.01, ***p<0.001.

**Figure S7. Serotonin does not change BNST-to-CeA low gamma (30-50 Hz) communication in either sex. (A1-A2)** Tone-evoked CeA and adBNST power spectra showing low gamma in ChR2-(grey) and ChR2+ (blue) female mice. **(A1)** In the CeA and **(A2)** BNST, ChR2+ (n=6-8) and ChR2-(n=5-9) females have equivalent levels of tone-evoked low gamma power. **(B)** Tone-evoked adBNST-CeA low gamma coherence was similar in ChR2-(grey, n=5) and ChR2+ (blue, n=6) female mice. **(C1-C2)** Tone-evoked CeA and adBNST power spectra showing low gamma in ChR2-(grey) and ChR2+ (blue) male mice. **(C1)** In the CeA and **(C2)** BNST, low gamma was similar in ChR2- (n=5-7) and ChR2+ (n=8-9) male mice. **(D)** There is no difference in tone-evoked BNST-CeA low gamma coherence between ChR2- (grey, n=5) and ChR2+ (blue, n=7) male mice. SEM shown in shaded regions. **(E-F)** Change in tone-evoked high gamma and low gamma power from pre-tone levels in the BNST. P-values under each group indicate whether the changed from zero was significant. **(E)** In females, tone-evoked high gamma power in the BNST decreased from pre-tone levels in ChR2-mice but not ChR2+ mice. In males, BNST high gamma power decreased in both genotypes. **(F)** In both sexes, tone-evoked low gamma power in the BNST decreased from pre-tone levels in both genotypes.

